# A biosensor to gauge protein homeostasis resilience differences in the nucleus compared to cytosol of mammalian cells

**DOI:** 10.1101/2021.04.19.440383

**Authors:** Candice B. Raeburn, Angelique Ormsby, Nagaraj S. Moily, Dezerae Cox, Simon Ebbinghaus, Alex Dickson, Gawain McColl, Danny M. Hatters

## Abstract

An extensive network of chaperones and other proteins maintain protein homeostasis and guard against inappropriate protein aggregation that is a hallmark of neurodegenerative diseases. Using a fluorescence resonance energy-based biosensor that simultaneously reports on intact cellular chaperone holdase activity and detrimental aggregation propensity, we investigated the buffering capacity of the systems managing protein homeostasis in the nucleus of the human cell line HEK293 compared to the cytosol. We found that the nucleus showed lower net holdase activity and reduced capacity to suppress protein aggregation, suggesting that the nuclear quality control resources are less effective compared to those in the cytosol. Aggregation of mutant huntingtin exon 1 protein (Httex1) in the cytosol appeared to deplete cytosolic chaperone supply by depleting holdase activity. Unexpectedly, the same stress increased holdase activity in the nucleus suggesting that proteostasis stress can trigger a rebalance of chaperone supply in different subcellular compartments. Collectively the findings suggest the cytosol has more capacity to manage imbalances in proteome foldedness than the nucleus, but chaperone supply can be redirected into the nucleus under conditions of proteostasis stress caused by cytosolic protein aggregation.

## Introduction

Protein homeostasis involves a network of quality control systems that ensures the proteome is properly translated, folded, delivered to the correct cellular location and turned over at appropriate times. When protein homeostasis becomes unbalanced, proteins become prone to misfolding leading to their mislocalisation and accumulation as aggregates (1,2). The imbalance of protein homeostasis is hypothesized to underlie the inappropriate protein misfolding and aggregation that arise in the brain of patients with common neurodegenerative diseases including Alzheimer’s, Parkinson’s, Huntington’s and motor neuron diseases (3,4). Tools that can measure proteostasis imbalance therefore offer capacity to explore the mechanisms involved.

Previously we developed a method that enabled a measure of the effectiveness of the quality control system in maintaining protein homeostasis (5). The method involved the use of a biosensor that comprised of a model protein that engages with protein quality control machinery such as chaperones. The biosensor reported on amount of the model protein bound to quality control proteins (which we call hereon call holdase activity) and on the ability of quality control proteins to repress inappropriate protein aggregation of the model protein. The model protein was a catalytically inactive variant of the prokaryotic RNAse barnase. Unfolded barnase is both permissive to aggregation and able to bind to Hsp70 and Hsp40 family chaperones (6). Barnase folding resembles a 2-state mechanism and the proportion of unfolded barnase relative to folded barnase can be predictably tuned by mutation (7). Hence a panel of barnase variants enable different biosensors tuned to different ratios of folded versus unfolded proteins. When the biosensor is expressed in cells the proportion of folded proteins can be measured by fluorescence resonance energy transfer (FRET) through N- and C- terminal fusions to fluorescent protein donors and acceptors (**Fig 1A**). We previously showed that the abundances of unfolded barnase is increased in cells relative to that predicted by analysis of purified proteins due to quality control machinery forming complexes with the unfolded-like conformations of the biosensor and partitioning it from the equilibrium of folding (6). Therefore, alterations in quality control levels influence the total abundance of unfolded-like barnase, which we can detect by FRET, and therefore determine changes in the overall quality control supply available as a measure of proteostasis capacity.

**Figure 1:**
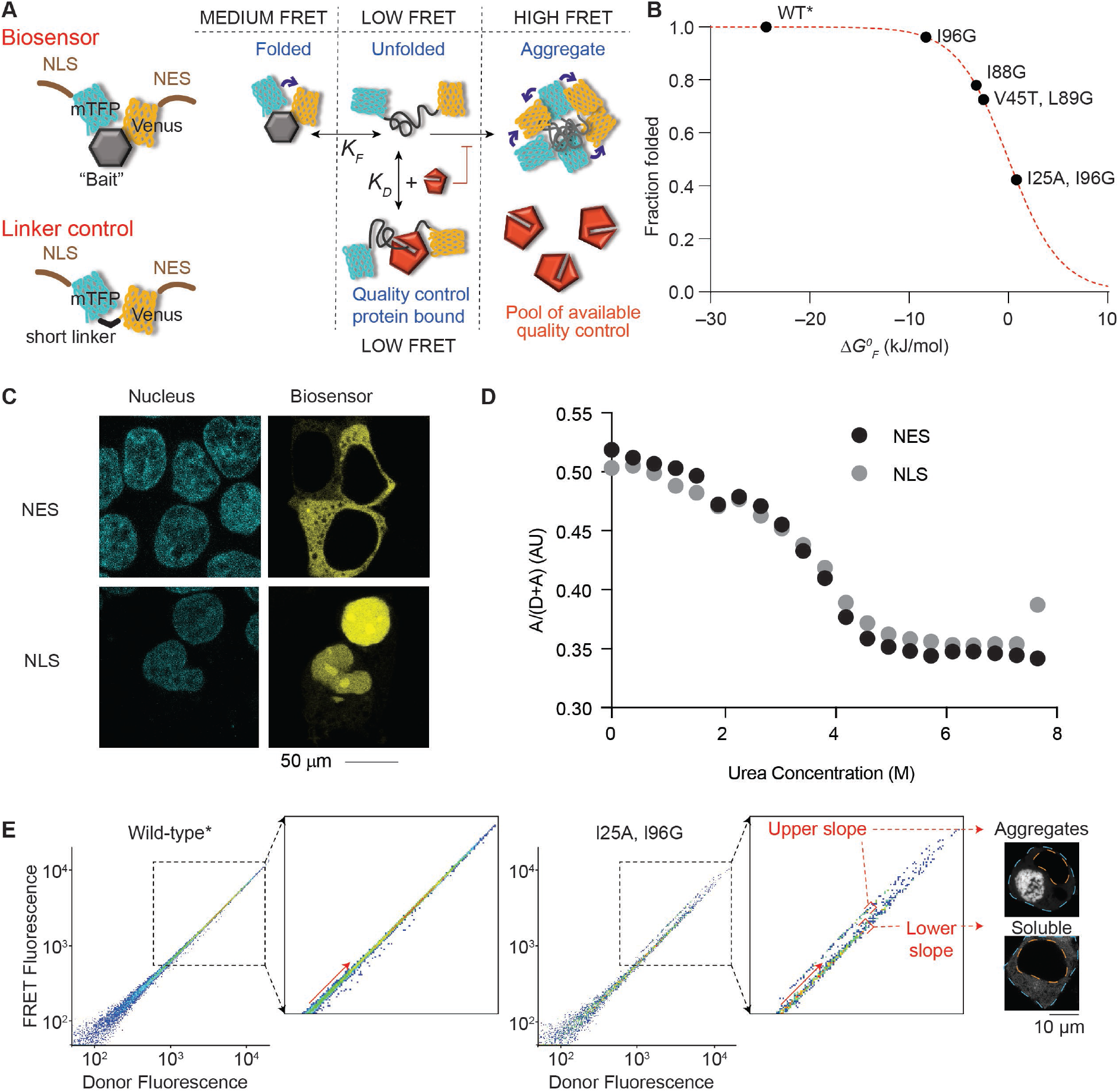
Targeting the barnase biosensor into the cytosol and nucleus. **A**. Schematic of how the biosensor works. The barnase protein is used as the “bait” for chaperones and is flanked with fluorescence proteins for Fluorescence Resonance Energy Transfer (FRET) measurements. A nuclear localization sequence (NLS) or nuclear export sequence (NES) is appended to the construct. The linker control has the barnase module omitted. **B**. Shown is the relationship between mutations in barnase, the effect on standard free energy of folding (Δ*Gº*_*F*_ at 20 °C), and predicted fraction folded for the various biosensor variants used in the study. Wild-type (WT) barnase is marked with * to denote it contains the catalytic inactivation mutation H102A. This mutation is present in all constructs in the study. **C**. Confocal images of HEK293T cells transiently transfected with either nuclear-or cytosol-targeting biosensor variant of the WT* barnase biosensor. The nucleus was visualized by Hoechst 33342 stain (cyan) and biosensor by Venus fluorescent protein fluorescence (yellow). **D**. Urea denaturation curves of WT* barnase biosensor variants as measured in cell lysates by FRET. **E**. Flow cytometry strategy for monitoring foldedness and aggregation. Here the donor and acceptor fluorescence of cells were measured by channels (FRET and Donor fluorescence was gated by the PE (575/25) and V500 (525/50) filters, respectively with the 405 nm laser). The inset highlights the changes that arise for cells bifurcated into “upper” and “lower” slope populations (division shown with red arrow). Representative cells collected from gates corresponding to the upper and lower slope populations imaged by confocal microscopy (grayscale). The orange dashed line denotes the nucleus boundary and the cyan dashed line the cell boundary.

The amount of biosensor aggregation can also provide a measure of overall chaperone activity. Our prior work devised a strategy that quantitatively measured aggregation as a complementary approach to foldedness (6). Here, we applied our biosensor system to examine how proteostasis balance is affected specifically within the nucleus and cytosol. We examine these local proteostasis changes that result from different triggers of stress either globally to the cell or locally within the cytoplasm or nucleus.

## Results

### Generation of nuclear targeted biosensors and validation of folding stabilities

The biosensor system comprises a suite of constructs whereby the barnase moiety has been mutated to display different standard free energies of folding (Δ*G*^*0*^_*F*_), which define the thermodynamic equilibrium of folding *K*_*f*_ (**Fig 1B**). These constructs contained a nuclear export sequence (NES), which leads to them being restricted to a cytosolic localization (6). To direct the constructs to the nucleus we removed the NES and fused a nuclear localization sequence (NLS) from the SV40 protein to the N-terminus (**Fig 1A; Table 1** for sequences used). The biosensor containing wild-type* (WT*) barnase (which is marked with * to denote it contains the catalytic inactivation mutation H102A that is used in all our constructs) was efficiently targeted into the nucleus (**Fig 1C**). All mutants of barnase showed a similar result (not shown).

**Table 1.**
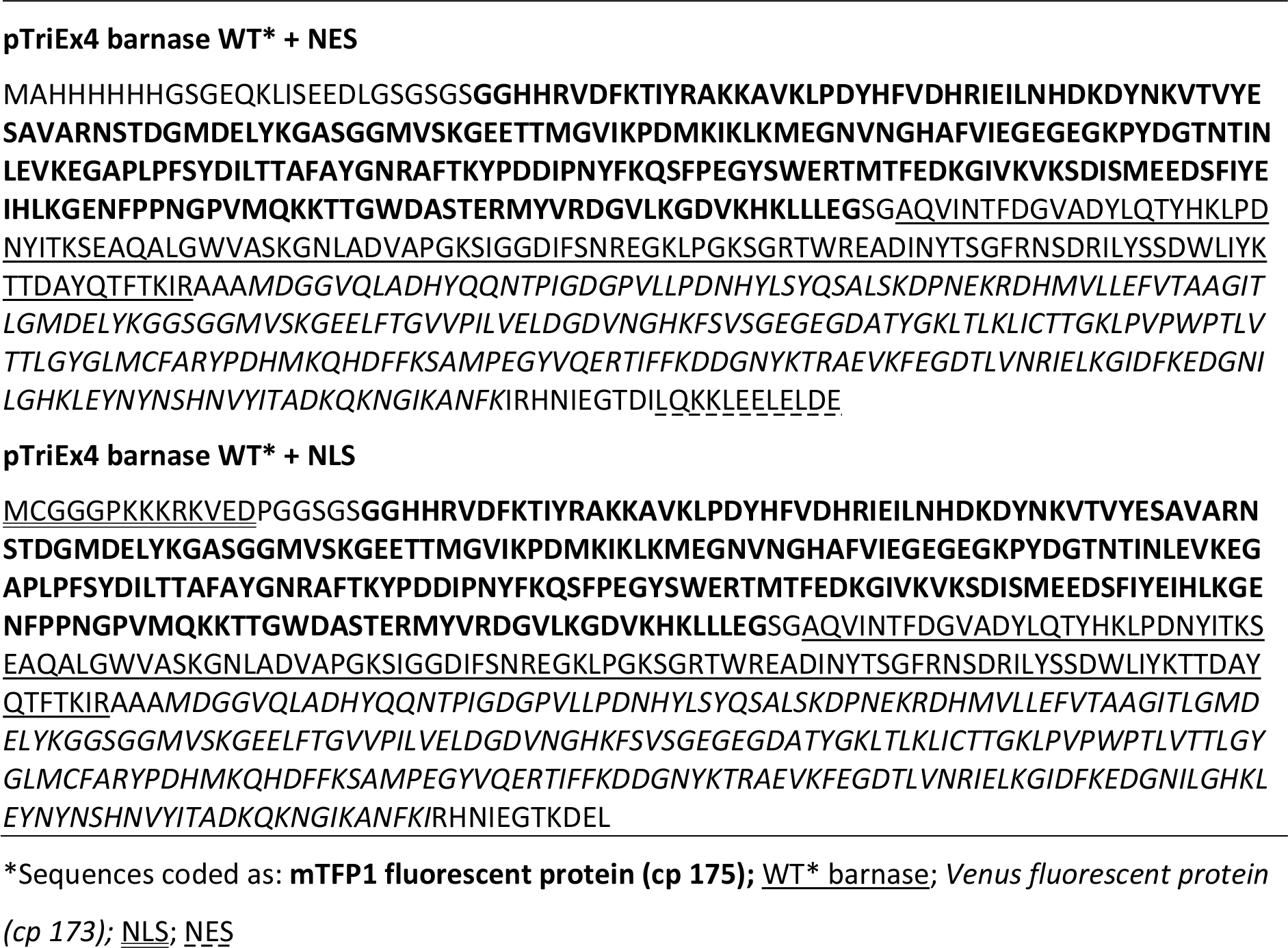
Sequence overview of the barnase biosensor constructs*.

Because the NLS and NES could themselves affect the folding equilibrium, we measured their effect on folding by a urea denaturation curve which showed no noticeable difference (**Fig 1D**). Hence, we concluded that the biosensors with NLS and NES are amenable to directly measure and compare protein homeostasis balances between the cytosol and nucleus.

The strategy to monitor both the abundances of unfolded barnase and aggregation behaviour involved a flow cytometry protocol we previously developed (6). In essence, cells expressing the biosensor bifurcate into two distinct FRET populations when cells are gated on acceptor fluorescence versus donor fluorescence. The donor fluorescence levels are proportional to expression level and FRET and both are approximately linearly dependent on the acceptor fluorescence. One of the populations comprises a “lower slope” FRET population that contains cells with only soluble barnase biosensor (i.e. a mixture of medium and low FRET states; **Fig 1A and E**), whereas the other contains an “upper slope” FRET population, which contains cells with aggregated biosensor (i.e. dominated by high FRET states) (6). The gradient of the lower slope population is proportional to the actual FRET value and hence informative to the abundance of unfolded barnase versus folded barnase (i.e. the average signal of low and medium FRET states from **Fig 1A**). We had previously shown that quality control machinery such as Hsp70 and Hsp40 proteins HSPA8 and DNAJB1, respectively, can bind to unfolded biosensor and hold it in an unfolded-like state that has a low FRET signal (6) as summarized conceptually in **Fig 1A**. This binding creates a pool of chaperone-bound biosensor that is partitioned from the equilibrium of folding. Greater partitioning leads to lower FRET signals, which can thus be used as a measure of the capacity of the quality control system to engage with the biosensor.

First, we examined whether the cytosolic and nuclear environments differentially affected the FRET readouts. This was achieved by expressing a FRET construct in which barnase was replaced with a short linker sequence that was not expected to be affected by changes in conformation or other ligand binding events. As such this linker should render the biosensor insensitive to folding-related effects and hence measures off-target influences on FRET signal as previously described (6). The linker control revealed a small (1.7%) but significant (*p* < 0.0001, Student’s t-test; 2-tailed) decrease in FRET in the nucleus compared to the cytosol between the NLS and NES tagged variants (**Fig 2A**). To correct for this influence, all the subsequent analyses involving the barnase biosensors were corrected for differences using the NLS and NES linker construct controls.

**Figure 2:**
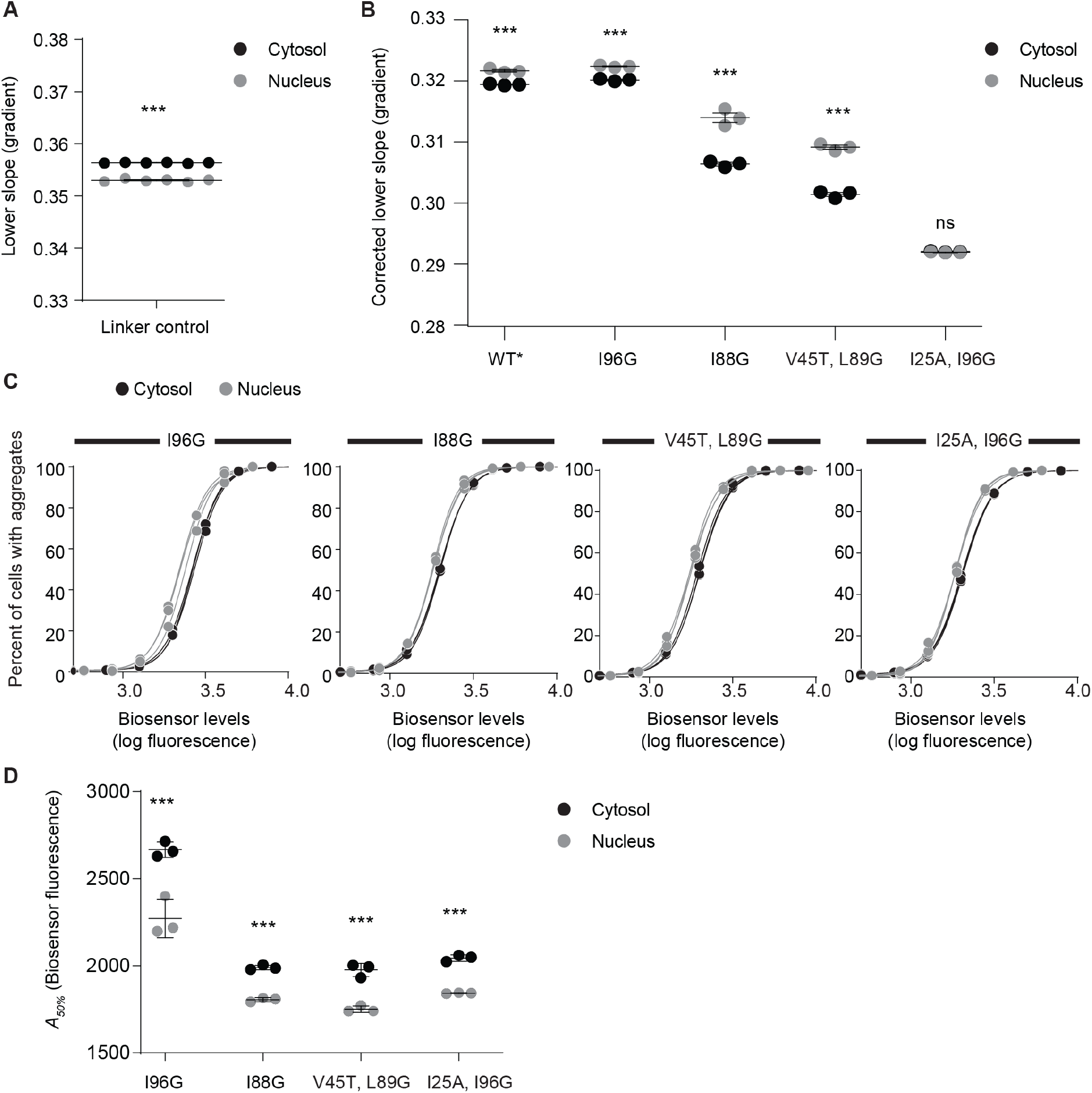
Reduced proteostasis resilience in the nucleus compared to the cytosol. All data in this figure relate to NLS or NES-tagged biosensor constructs transfected in HEK293 cells and analysed by flow cytometry. Individual biological replicates shown with means ± S.D. and with differences (nucleus v cytosol) assessed by Student’s t-test (2-tailed); *** p < 0.001, ns => 0.05. **A**. Effect of intrinsic FRET differences in nucleus versus cytosol assessed with the linker control. **B**. Analysis of different barnase mutations. All data were corrected for background effects using the linker control. **C**. Assessment of aggregation. Shown are cells binned into different biosensor levels (based on Venus fluorescence) and each bin assessed for percent in upper slope versus lower slope populations. Curves are fits to Hill equation. Data were fitted independently within each replicate dataset (n=3). **D**. Shown are biosensor concentrations at which 50% of cells have aggregates (*A*_*50%*_), derived from the Hill equation fits shown in panel C.

Next, we examined the effectiveness of protein quality control systems to interact with the biosensor in the nucleus versus the cytosol. For this we examined four previously characterized mutants of barnase in addition to wild-type* that have variable Δ*G*^*0*^_*F*_ values and therefore different proportions of folded to unfolded barnase at equilibrium (**Fig 1B**). After correction for off-target FRET changes with the linker control, all nuclear-targeted biosensor variants had overall significantly higher FRET values for the lower slope populations than the cytosol-targeted biosensors, except for the most destabilised variant (I25A I96G), which is predicted to be substantially unfolded and therefore possibly outside the dynamic range that can be detected (**Fig 2B**). The higher FRET values in the nucleus are therefore indicative of less unfolded-like barnase conformations being held in complex with chaperones that would otherwise be partitioned from the equilibrium of folding. The results therefore suggested that the pool of chaperones that can bind to the biosensor is lower in the nucleus than the cytosol.

To examine the aggregation propensity of the barnase biosensors, we applied our previously devised method of determining the concentration of barnase at which 50% of the cells contain aggregates (*A*_*50%*_) (6). *A*_*50%*_ values are derived from plots of the proportions of cells partitioning in the upper slope for a given expression level of barnase in cells (**Fig 2C-D**). Consistent with prior findings (6), WT* barnase did not aggregate and the less stable mutants (i.e. those with higher Δ*G*^*0*^_*F*_ values) were more sensitive to aggregation as determined by lower *A*_*50%*_ values (**Fig 2D**). For all the barnase variants that aggregated, the *A*_*50%*_ values were lower in the nucleus compared to the cytosol (**Fig 2C-D**). These results indicated that barnase is inherently more aggregation prone in the nucleus than the cytosol, and therefore strengthens the conclusion that there are less quality control proteins in the nucleus that are able to bind to and stabilize barnase and prevent aggregation.

### Hsp70 and Hsp40 chaperone systems more robustly mitigate unfolded proteins from aggregating in the cytosol than the nucleus

Hsp70 isoforms HSPA1B and HSPA8 were previously found in immunoprecipitation experiments as major chaperone interactors to the destabilized barnase mutants (6). We found that related Hsp70 family member HSPA1A could modulate both the amount of unfolded-like barnase and the amount of barnase aggregation, suggesting it could bind to and stabilize an unfolded-like conformation of barnase (6). While these Hsp70 isoforms are highly abundant in the cytosol (8), it was unclear as to how modulating their supply might propagate changes in the nucleus or cytosol.

To examine this question, we co-transfected HSPA1A and a specific Hsp40 cofactor DNAJB1 (9) with the barnase biosensors and analysed the cells after 48 hours culture. The transfected HSPA1A and DNAJB1 showed a mostly cytosolic enrichment (**Fig 3A**). The co-transfected chaperones significantly reduced the biosensor lower slope gradients in the cytosol (**Fig 3B**), consistent with a greater abundance of chaperone bound to unfolded barnase. The treatments also increased the *A*_*50%*_ values indicating that the chaperones effectively suppressed inappropriate aggregation (**Fig 3C**). However, these effects appeared more muted in the nucleus indicating that chaperone overexpression preferentially deepens the pool of chaperone supply in the cytosol, which likely is explained by the transfected chaperone being mostly restricted to the cytosol.

**Figure 3:**
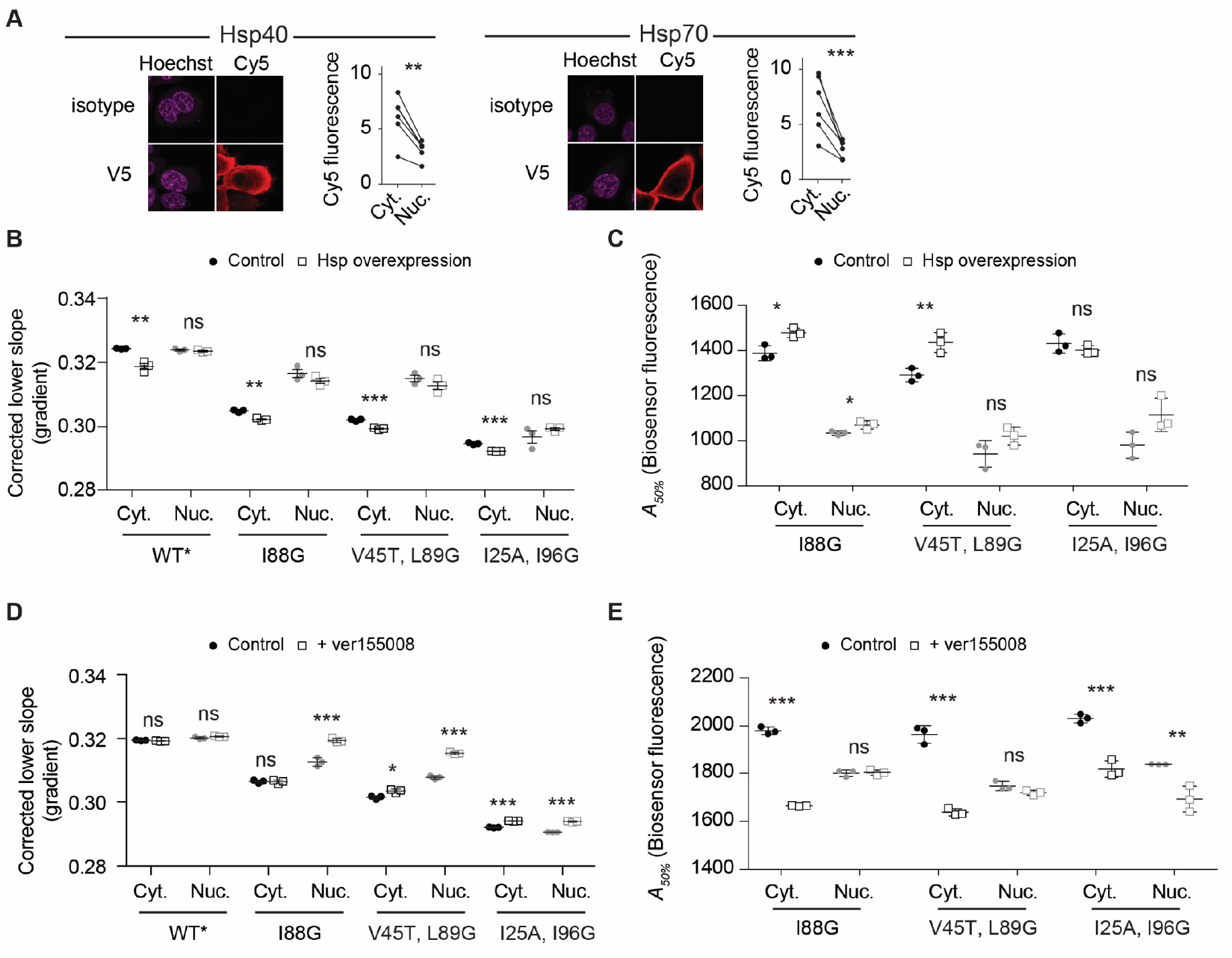
Cytosolic Hsp70 and Hsp40 activity provide depth in proteostasis resistance against protein misfolding and aggregation. **A**. Shown are immunofluorescence micrographs of HEK293T cells transiently transfected with either V5-tagged Hsp40 or Hsp70 proteins (DNAJB1 and HSPA1A respectively). The nucleus is stained with Hoechst 33342 and chaperone with Cy5 labelled anti-V5 antibody (or isotype control for specificity). Graphs indicate quantitation with paired Student’s t-test results shown (2-tailed, paired); *** p < 0.001, ** p < 0.01. Data points represent immunofluorescence intensity in single cells (paired by mean cytosol and mean nucleus). **B**. Lower slope analysis by flow cytometry of HEK293 cells co transfected with the biosensors, DNAJB1 and HSPA1A or control (a non-fluorescent derivative of GFP (Y66L Emerald (13)) for 48 hours. Data points indicate biological replicates, bars indicate means ± S.D. Student’s t-test results are shown (2-tailed; control v overexpression); *** p < 0.001, ** p < 0.01, * p < 0.05, ns => 0.05. **C**. Aggregation analysis (*A*_*50%*_) using the same treatments and conditions as for panel B. **D**. Lower slope analysis of HEK293 cells after transfection with the biosensors for 18 hours and a further treatment with 20 μM Hsp70 inhibitor VER-155008 for 18 h (versus vehicle control). Data is presented as per panel B. **E**. Aggregation analysis as presented for the other panels above.

To further probe the role of Hsp70 activity we inhibited Hsp70 on cells (without overexpressed chaperones) using the small molecule inhibitor VER-155008, which competitively binds to the ATP-binding pocket of Hsp70 family proteins and impairs substrate binding (10). This treatment increased the FRET values of the lower slope populations in the nucleus but not the cytosol (**Fig 3D**). This result suggested that while Hsp40 and 70 proteins are more effective at binding barnase in the cytosol, there was higher redundancy and flexibility in the cytosol to absorb a reduced Hsp70 activity than in the nucleus. Hence the network appeared more vulnerable to collapse in the nucleus upon stresses to proteostasis systems. However, the increased sensitivity to proteostasis imbalance in the nucleus was not seen in terms of aggregation. Indeed, aggregation of the biosensor was far more disproportionately enhanced in the cytosol than the nucleus (**Fig 3E**). These findings suggested that the correlation of holdase activity and aggregation can be decoupled when specific elements of the proteostasis network are impaired and that this effect may arise through redundant holdase activity in the cytosol from non-Hsp70 chaperones that are overall less effective at preventing aggregation than Hsp70.

### Aggregation of mutant Htt exon 1 in the cytosol propagates proteostasis imbalances in the cytosol and nucleus

Next we investigated the quality control supply in the nucleus and cytosol in the context of disease-related protein misfolding and aggregation. For this we co-expressed the biosensors with mutant Huntington exon 1 fragment containing 97 glutamines in the polyglutamine repeat sequence (Httex1_97Q_) fused to GFP, which forms cytosolic perinuclear inclusions in HEK293T cells (11). Mutant Httex1 fragments with polyglutamine sequences longer than 36 glutamines accumulate into inclusion bodies in neurons of Huntington Disease patients and have been implicated to direct a maladaptation in protein quality control (12). Because the fluorescence of GFP interferes with our capacity to monitor FRET, we used a variant of GFP that is non-fluorescent as characterized previously (13) and also added a 6 amino acid tetracysteine motif for post hoc detection by ReAsH biarsenical dye binding (14). Live cells were examined for inclusions, which were detectable as spherical pearl-like structures under transmission imaging of confocal microscopy, imaged for FRET and then post hoc analyzed by ReAsH staining to validate the inclusion structure. Because we needed to fix the cells after imaging, which reduces ReAsH staining, we were not able to ascertain cells containing only diffuse cytosolic Httex1_97Q_. Using this approach, we observed the barnase biosensor as enriched at the periphery of the inclusions suggesting a degree of co-aggregation or co-recruitment to the inclusions (**Fig 4A**). This was both true for the WT* barnase biosensor, which does not aggregate by its own volition, and for the nucleus-targeted biosensors suggesting that the biosensors were kinetically trapped on the surface of the Httex1 inclusion. To assess whether the biosensor was self-aggregated at the molecular scale we determined their FRET using a ratiometric analysis of the fluorescence (**Fig 4A**). Indeed, the biosensor enriched at the inclusion periphery appeared to have higher FRET than when more distal from the inclusions in either the nucleus or cytosol. We further assessed the aggregation state using fluorescence recovery after photobleaching (FRAP) (**Fig 4B**). A small section of the biosensor was targeted for bleaching on the periphery of the Httex1 inclusion. Both nucleus and cytosol targeted biosensors showed little to no recovery after bleaching, indicating the protein was in an immobile state on the seconds timescale (**Fig 4C**).

**Figure 4:**
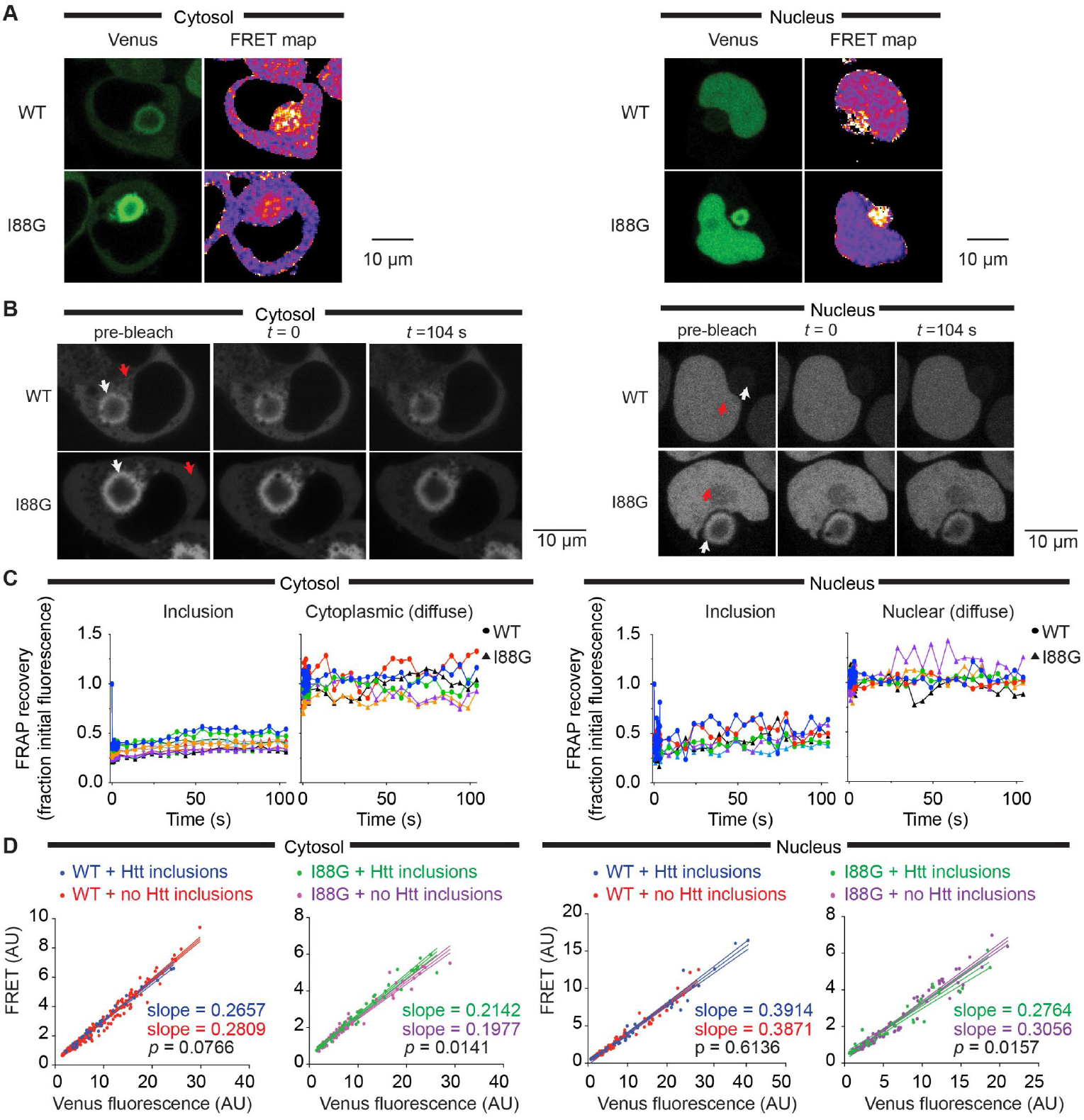
Huntingtin exon 1 aggregation in the cytosol manifests proteostasis imbalances in the nucleus and cytoplasm. **A**. Confocal micrographs showing the values proportional to the fluorescence ratio of acceptor/donor (Venus/mTFP) of the biosensors co-expressed with mutant Httex1_97Q_ fused to a non-fluorescent mutant of GFP. Selected cells are those with Httex1 inclusions, identified post-hoc as described in the methods. Nuclear targeted biosensors are on the right and cytosolic targeted biosensors on the left (same format for each panel). The scale of the FRET map is colour coded from blue to magenta to yellow corresponding to lowest to highest FRET. **B**. Fluorescence recovery after photobleaching (FRAP) of biosensor at the periphery of the Httex1_97Q_ inclusion. Arrows indicate region of bleaching. **C**. Quantitation of the data in panel B, tracking the recovery of fluorescence in the bleached zone. Each colour depicts the time course of an individual cell. **D**. Confocal microscopy FRET fluorescence values within cells distal to the inclusion. FRET fluorescence was measured by exciting at 458nm (mTFP1 excitation) and collecting the emission at 510-560 nm (Venus emission). Each dot represents the average fluorescence derived from a single cell value. Solid lines show line of best fit from a linear regression with dashed lines showing 95% confidence intervals. P-value was determined by two-tailed t-test.

Because the aggregation of the biosensor at the periphery of the inclusion is a confounding factor in whole cell fluorescence analysis by our flow cytometry methods, we instead measured FRET by microscopy targeting small subregions of the cells away from the inclusions. This analysis revealed that the presence of Httex1_97Q_ inclusions had no bearing on the FRET of the WT* barnase biosensor outside that associated with the inclusion periphery (**Fig 4D**). By contrast, the I88G barnase biosensor showed significantly changed FRET values regions outside the Httex1_97Q_ inclusions compared to cells without inclusions at all. In the case of the I88G barnase biosensor targeted to the cytosol, the FRET was increased, which suggested a reduced overall holdase activity of chaperones in the cytosol arising from Httex1_97Q_ aggregation. However, in the case of the I88G barnase biosensor targeted to the nucleus the FRET was decreased. This result is consistent with an elevated holdase activity in the nucleus when Httex1_97Q_ aggregates form. This result therefore suggests when protein aggregation occurs in the cytosol that cells can move the pool of quality control machinery from the cytosol into the nucleus as part of a global coordinated stress response.

## Discussion

Our studies show that the balance of resources required to manage proteostasis is different in the cytosol relative to the nucleus. We find evidence for there being a lower supply of chaperone capacity in the nucleus that is able to bind to the unfolded barnase and prevent its aggregation. When we supplemented the cells with additional Hsp70 and Hsp40 protein by their overexpression (HSPA1A and DNAJB1), we increased holdase activity in the cytosol and lowered the aggregation potential consistent with these chaperones exerting a critical activity to bind unfolded proteins and prevent their aggregation. When we pharmacologically inhibited the Hsp70 chaperone system we observed a disproportionate impact on aggregation in the cytosol, concordant with the cytosol being more richly dependent on Hsp40 and Hsp70 chaperone-based networks to prevent protein aggregation.

Overall, these results are consistent with the greater requirement of the Hsp70 chaperone system to engage with unfolded or aggregation-prone proteins in the cytosol. This finding is consistent with the high abundance of these chaperones in the cytosol (8), which is not surprising given that most proteins are synthesized in the cytosol or endoplasmic reticulum.

Our findings with mutant Httex1_97Q_ indicate that the aggregation in the cytosol can manifest dysfunction in quality control capacity in both the cytosol and nucleus. Consistent with prior findings that protein aggregation can sequester quality control resources away from “housekeeping” activities and lead to metastably-folded proteins aggregating (15), we found that the pool of resources binding to unfolded barnase biosensor decreased in the cytosol. Prior studies have found that Hsp70 and Hsp40 proteins are recruited into inclusions formed by Httex1_97Q_ and similar proteins with expanded polyglutamine sequences (16-19). One function for this recruitment may be disaggregation, in light of recent findings showing the Hsp70 based chaperone machinery can dissociate amyloid fibrils (20). More unexpected however was the finding that there was an increase in unfolded barnase in the nucleus, which suggests that chaperones are redirected into the nucleus under stress. Hsp70 is known to translocate from the cytosol into the nucleus upon heat shock (21-23), suggesting there is a dynamic capacity for quality control machinery activity in the nucleus under times of stress. This translocation is regulated by the Hikeshi nuclear import carrier, which is crucial for cells to recover from heat shock stress (24). DNAJB1 can also deliver misfolded protein into the nucleus for degradation (25).

The other notable result from our study was the recruitment of WT* barnase biosensor to the Httex1_97Q_ inclusion. We have never observed the wild-type* biosensor to aggregate when expressed on its own suggesting that the inclusion provides a mechanism to recruit this protein to the surface. One interesting possibility is that a small fraction of the biosensor remains in complex with chaperones; and that these complexes are recruited to the surface of the inclusion by Hsp70 -based triage mechanisms that more generally handle misfolded proteins in the cell. The different extent of WT* biosensor foldedness in the nucleus compared to the cytoplasm supports the conclusion that some of the wild-type barnase is partitioned from the equilibrium of folding in an unfolded-like state. Discrete bodies containing misfolded protein including the JUNQ, aggresome and Q-bodies have been proposed as cellular depots for processing protein aggregates, and are enriched with different Hsp70 and Hsp40 proteins (26,27). In addition, Hsp70 has been proposed to engage with the surface of protein aggregates to act as a disaggregase (20). Hence, the capture of wild-type* biosensor may be indicative of a wider network of chaperone client interactions, protein aggregate bodies in the cell and a broader interconnected quality control network. And thus chaperones may have a broader function as a kind of lubricant constantly interfacing with unfolded proteins and aggregating proteins.

## Materials and Methods

### Expression constructs

The cytosolic FRET barnase biosensor library expressed in the pTriEx4 vector were prepared as previously described (5). In brief, the barnase moiety was flanked by circularly permuted mTFP1 cp175 and Venus cp173 fluorescent proteins. Nuclear localised FRET barnase was generated by the addition of a N-terminal SV40 NLS sequence to the original cytosolic barnase using a synthesized gene cassette containing the relevant localization sequences (GeneArt (Thermofisher), Waltham, Massachusetts) and standard restriction endonuclease-based ligation methods. For generation of individual mutants of targeting biosensor, the WT* barnase biosensor kernel was replaced by the barnase mutant of choice. This was achieved by double-digestion of both the desired barnase mutant and nuclear targeting construct plasmids at BamHI and KpnI restriction sites. The tetracysteine tagged Httex1 construct containing a tetracysteine tag at the C-terminus of the Httex1 (TC1 (28)), and a non-fluorescent mutant of Emerald fluorescent protein (Em), Y66L (13), was generated in-house to yield a plasmid named Httex1_97Q_TC1-Em Y66L in the pT-Rex vector (Invitrogen). The pT-Rex Em Y66L construct alone was also generated in-house as described previously (13). V5-tagged chaperone proteins were overexpressed from pcDNA5/FRT/TO V5 DNAJB1 and pcDNA5/FRT/TO V5 HSPA1A provided as gifts from Harm Kampinga (29).via Addgene, Watertown, Massachusetts.

### Cell culture

HEK293T cells were maintained in DMEM supplemented with 10% (w/v) fetal calf serum and 1 mM glutamine in a 37°C humidified incubator with 5% v/v atmospheric CO_2_. Cells were seeded in poly-L-lysine coated plates. For microscopy experiments cells were plated at 3 × 10^5^ cells/ml in an 8 well µ-slide (Ibidi, Martinsreid, Germany). For flow cytometry experiments cells were seeded at 1.1 × 10^5^ cells/ml in a 48 well plate. Cells were transiently transfected with Lipofectamine 3000 reagent as per manufacturer’s instructions (Life Technologies, Thermofisher). For Barnase and Httex1 co-transfections, the transfection was done in a way to decouple the expression of the two plasmids.

HSP70 was inhibited with 20 µM VER-155008 (cat #SML0271, Sigma-Aldrich, St. Louis, Missouri) in culture media for 18 h.

### Microscopy

Cells were imaged on a TCS SP5 confocal microscope (Leica Biosystems, Nussloch, Germany). For immunofluorescence, cells were fixed in 4% w/v paraformaldehyde for 15 mins at room temperature, washed with phosphate buffered saline (PBS), and permeabilized in 0.5% v/v Triton X-100 in PBS (Sigma-Aldrich) for 30 mins. After incubation in blocking solution (5% w/v bovine serum albumin in 0.3% v/v Triton X-100 in PBS), cells were incubated with anti-V5 (1:250 dilution in 1% w/v bovine serum albumin in 0.3% v/v Triton X-100 in PBS) (cat #ab27671, Abcam, Cambridge, United Kingdom) overnight at 4°C. Cells were then washed in 1% w/v ovine serum albumin in 0.3% v/v Triton X-100 in PBS before being stained with anti-mouse cy5 (1:500 dilution in PBS) for 30 min at room temperature. Prior to confocal imaging, cells were stained with Hoechst 33342 (ThermoFisher).

### Image analysis

Confocal images were analysed using custom analysis scripts for FIJI (30) and Python (v 3.6.7), available alongside example datasets at doi.org/10.5281/zenodo.4686851.

In the case of immunofluorescence measurements, whole cells and nuclei were initially identified using the machine learning package CellPose (31). Segmentation was performed on the Cy5-labelled anti-V5 antibody and Hoechst channels (633 nm excitation, 650-750 nm emission and 405 nm excitation, 410-450 nm emission respectively) to identify the whole cell and nuclei regions of interest (ROI) respectively. Per-pixel information for each ROI was then collected and the nuclei ROIs removed from the whole cell to yield the cytosolic population. Finally, the fluorescence intensity for the nucleus and cytosol was determined from the mean of all pixels in each compartment.

To quantify fluorescence recovery after photobleaching (FRAP), ROI for individual bleach spots were defined via automatic Otsu thresholding of the first bleaching frame. Identical ROI’s were then manually placed for the adjacent (non-bleached) and background regions. Where necessary, ROI positions were manually adjusted across timepoints to account for cellular drift. The mean intensity was calculated for each ROI, and both bleached and non-bleached ROIs were then corrected against the corresponding background ROI for each time point, generating *B*_*corr*_ and *NB*_*corr*_ respectively. The ratio of *B*_*corr*_ / *NB*_*corr*_ at each time point was finally normalised to the pre-bleach ratio of *B*_*corr*_ / *NB*_*corr*_ to yield the relative recovery.

In the case of FRET measurements, whole cells, nuclei and Httex1 inclusions were initially identified using CellPose (31) as described above. In this case, segmentation was performed on the acceptor channel (488 nm excitation, 510–560 nm emission), computationally inverted acceptor channel and ReAsH channel (561 nm excitation, 610–680 nm emission) for whole cells, nuclei and inclusions respectively. After manual inspection to ensure the accuracy of each round of segmentation, per-pixel intensity values for each ROI were collected. In the case of cytosolic barnase variants, both the nuclei and aggregate features were excluded from the whole cell to yield the diffuse barnase population. In the case of nuclear barnase variants, any aggregate ROI within the nuclei ROI were similarly excluded to yield the diffuse barnase population. Finally, the relative FRET for each ROI was calculated as the mean per-pixel intensity in the FRET channel (458 nm excitation, 510-560 nm emission).

### Flow cytometry

After 24 h (drug treatments) or 48 h (co-transfections) post-transfection, cells were washed and harvested by gentle pipetting in PBS. Cells were analysed via flow cytometry as described previously (6). In short, 150 µl of cell suspension was analysed at flowrate of 3 μl/sec on a BD LSRFortessa cell analyser (BD Biosciences, North Ryde, NSW, Australia). Acceptor (Venus) fluorescence was collected with the 488 nm laser and FITC (530/30) filter. Acceptor sensitized emission (FRET) and donor (mTFP1) fluorescence were collected with the 405 nm laser with PE (575/25) and V500 (525/50) filters, respectively. All flow cytometry data were processed with FlowJo (version 10, Tree Star Inc, Ashland, Oregon) to exclude cell debris, cell aggregates and untransfected cells. The Venus channel was compensated to remove bleed through from mTFP1 and FRET channels using donor only. Data were analysed in MATLAB (version 9, MathWorks, Natick, Massachusetts). The gating strategy and associated data analysis protocols are detailed previously (32).

### Urea denaturation curves

Urea denaturation curves were measured on cell lysates expressing the biosensors in 96 well format. In essence, 80 µl of samples were prepared containing 0 M to 8 M urea in PBS. Lysates were prepared from cells 24 h after transfection (wild-type* with NES and NLS tags) by pipetting in 20 mM Tris pH 8.0, 2 mM MgCl_2_, 1% v/v Triton X-100, 1 × EDTA-free protease inhibitor (Roche, Basel, Switzerland), 150 mM NaCl, 20 U ml^−1^ benzonase, 1 mM PMSF. Aggregates and cell debris were pelleted by centrifugation at 16,000 *g* for 10 min at 4 °C. 5 µl supernatant was added to each urea concentration. As the measurements were ratiometric and both fluorophores were on the same molecule, samples were not matched for protein concentration. Fluorescence readings (430 nm excitation, 492 nm emission and 532 nm emission) were measured at 23 °C using a Clariostar microplate reader (BMG Labtech, Mornington, Victoria, Australia) every 15 min for 4 h. Relative FRET efficiencies (calculated as Acceptor fluorescence/[Donor fluorescence + Acceptor fluorescence]) were averaged across readings and fit to a two-state unfolding model as described previously (6).

## Notes

https://zenodo.org/record/4686851#.YH0nW-gzaiM

### Competing Interest Statement

The authors have declared no competing interest.

